# A simplified hybrid capture approach retains high specificity and enables PCR-free workflow

**DOI:** 10.1101/2025.03.18.644033

**Authors:** Adeline Huizhen Mah, Xiaodong Qi, Junhua Zhao, Kelly Wiseman, Laure Edoli, Kyle Metcalfe, Kyle Donohoe, Micah Ojeda, Sophie Billings, Juan Moreno, Marina McCowin, Ben Krajacich, Kevin Green, Ramkrishna Adhikary, Andrew Boddicker, Joshua Chan, Peter Mains, Bryan Lajoie, Sean Devitt, Semyon Kruglyak, Shawn Levy, Michael Previte

**Affiliations:** Element Biosciences

**Author notes:** equal contribution (Adeline Huizhen Mah, Xiaodong Qi, Junhua Zhao).

## Abstract

Hybrid capture is a critical technology for selective enrichment of genomic regions of interest, enabling cost-effective focused sequencing in both clinical and research applications. We present a simplified hybrid capture approach that eliminates complexities typically associated with hybridization-based selection methods by directly loading the hybridization product onto the sequencing flow cell. The workflow removes bead-based steps, washes, and post-hybridization PCR, while retaining high capture specificity and library complexity. The approach is enabled by the development of a streptavidin flow cell surface, a method to circularize and amplify captured targets on the flow cell, and a fast hybridization protocol. We demonstrate application across targeted panel sizes and improvements to library complexity and variant calling. We also show how the approach can be used to create an entirely PCR-free targeted sequencing workflow that further improves variant calling and enables the detection of repeat expansions.

## Introduction

Hybridization-based target selection, also known as hybrid capture, is a powerful molecular biology technique. It enables researchers to selectively enrich specific regions of the genome for high-throughput sequencing, facilitating deeper analysis and understanding of targeted loci. In this technique, DNA or RNA probes (baits) are designed to be complementary to the regions of interest in the genome. These baits are then used to capture and enrich target sequences from a fragmented genomic DNA library. The process generally involves hybridizing the library fragments to the baits, capturing the baits with hybridized library, followed by washing away unbound fragments. The captured target sequences are then eluted or directly amplified for downstream sequencing.

Hybridization based methods are nearly ubiquitous in genomics, with applications in basic and translational research^1-12^. The solution-phase hybrid capture method is the primary approach used today, driven by optimization of hybridization-based probe designs and the broad availability of targeted panels^2,3,5,8,9,11^. A popular application is exome sequencing, which focuses on the protein-coding regions of the genome that harbor the majority of known disease-causing mutations while comprising only ∼1% of the human genome. The approach is also applied to smaller panels allowing researchers to focus on specific genes or genomic regions of interest across applications such as cancer genomics, microbial genomics, forensic science and agriculture^12-19^. Hybrid capture offers several advantages over alternative target enrichment methods such as multiplex PCR. Most notably, hybrid capture can be applied to a wide range of target sizes, from a few kilobases to many megabases that cover the entire exome^9^. The method provides cost-savings in sequencing while delivering relatively uniform coverage across the target regions, minimizing biases, and ensuring accurate representation of the sequences^10^.

A major drawback of targeted hybridization methods is a lengthy and complex workflow. The post-hybridization steps of the workflow have remained largely the same since their original development with minor optimizations in hybridization time and the use of refined buffers and wash steps to increase specificity. The methods nearly universally use magnetic beads containing streptavidin to bind the biotinylated oligo baits that have been hybridized to the target library.

Following bead binding, a series of temperature-controlled washes to remove the unbound and non-specific material are performed. These steps are essential in achieving a high on-target rate, but they lead to a large loss of DNA, thus requiring PCR amplification of the captured DNA to generate a final library with sufficient input material for sequencing. The common use of two rounds of PCR throughout the hybridization selection process followed by exacting washes and the need to retain bead materials through multiple wash steps make the post-hybridization steps a key source of variability in the assay. The post-hybridization PCR also leads to reduced library complexity through a sampling effect, where random amplification of DNA molecules results in the loss of some rare sequences. Streamlining this workflow would substantially reduce turnaround time and increase efficiency across sequencing applications.

Here we describe Trinity, a modified hybrid capture workflow, that addresses the workflow challenges of the traditional approach, while retaining its key advantages and extending its capabilities. These changes eliminate the need for post-hybridization PCR amplification, multiple wash steps, and the use of streptavidin beads. Traditional target enrichment workflows require 12-24 hours to complete^8,9,17^. With the Trinity workflow, the entire process from the start of library preparation to sequencer loading can be completed in as fast as 5 hours. The sequencing results demonstrate reduced duplicate rates, improved on-target rates in smaller panels, and higher accuracy indel calling. When combined with a PCR-free library prep, Trinity also supports an entirely PCR-free targeted sequencing workflow.

## Results

### Workflow and performance comparison

Figure 1A illustrates the traditional hybrid capture workflow as originally described in Gnirke et al^8^. We set out to eliminate all the manual post-hybridization steps by enabling direct loading of the hybridization product onto the flow cell. A major challenge with this approach is that the hybridization product contains not only library fragments bound to the baits (i.e. the targeted regions) but also the unbound library fragments, which have far greater abundance. To overcome this specificity challenge, we developed a passivated streptavidin flow cell surface that could directly capture the biotinylated baits with bound library elements, without capturing the unbound (off-target) library elements. This flow cell surface obviates the need for the streptavidin beads and temperature-controlled bead washing steps that require centrifugation and often multiple subsequent washes^20^. In the traditional workflow, off-target sequences are inadvertently captured through two mechanisms: (1) hybridization between repetitive elements within the genomic inserts themselves, and (2) adapter-mediated cross-hybridization, where complementary adapter sequences anneal between on-target and off-target library fragments.

**Figure 1.**
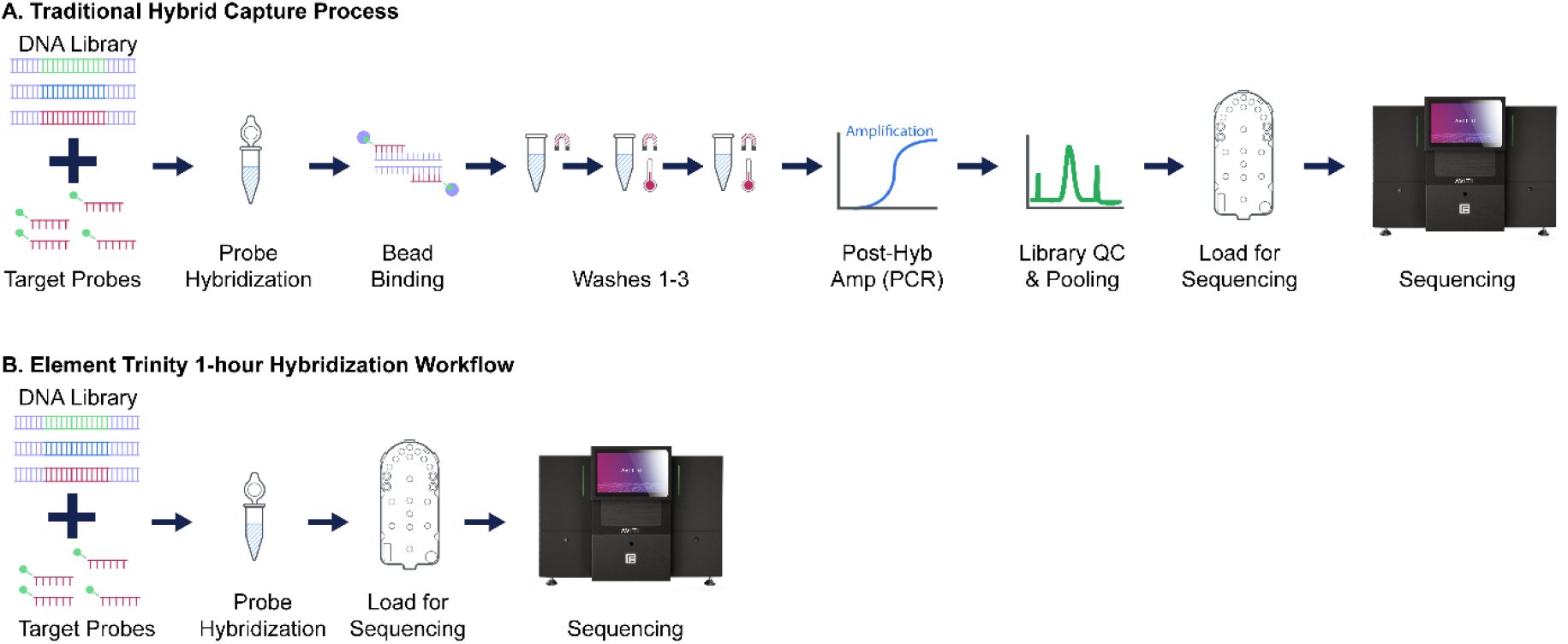
Comparison of the traditional hybrid selection workflow (A) and Trinity (B). The simplified Trinity workflow eliminates bead binding, washes, post-hyb PCR, and library QC.

These are two major contributors to off-target sequences in the hybrid capture workflow^21^. The insert annealing is usually prevented by adding repetitive blocking DNA^22^. With our new flow cell, biotinylated probe/library complexes are captured on the surface, and we perform an on-flowcell circularization followed by rolling circle amplification^23^. Our use of a circularization oligonucleotide prevents adapter annealing formation and results in high on-target performance, without the need for adapter-specific “blocker” reagents that are typically used to achieve high specificity^24^. The RCA product is directly sequenced on the AVITI instrument as previously described^23^. Figure 1B illustrates the Trinity workflow. The initial steps leading to hybridization remain unchanged, but the workflows diverge significantly thereafter, with Trinity proceeding to sequencing following hybridization. In addition to eliminating the bead capture, wash steps, post-hybridization PCR, and QC, we optimized conditions to shorten the hybridization time to one hour while maintaining high specificity and uniformity. The optimization enabled us to go from library preparation to loading of the sequencer within a single work shift. Specifically, we completed the workflow in under 5 hours with 2.5 hours for the library preparation, 1 hour for pooling and pre-hybridization steps, 1 hour for the hybridization, and 20 minutes for setting up the sequencing run. For the traditional workflow, the additional steps required us to break the workflow into two days.

To evaluate the performance of the Trinity assay, we compared the traditional workflow with a 16-hour hybridization to the Trinity workflow with a 1-hour hybridization across two bait set vendors. Table 1 shows the average results for 24 samples in each workflow. Trinity performance exceeded the traditional approach across most metrics including fold-80, duplicate rate, and variant calling accuracy – particularly for indels. The 16-hour traditional approach from vendor B showed a higher on-target rate, but this was partially offset by the higher duplicate rate. Additionally, Trinity demonstrated higher variant calling accuracy when starting with the same number of input reads.

**Table 1:** Metric comparison of traditional exome approach with 16-hour hybridization and Trinity 1-hour hybridization across two bait set manufacturers. Vendor A Sequenced with 2×150 reads and Vendor B sequenced with 2×75 reads. Vendor panels differ in size and content. All analysis performed with 30 million reads per sample and metrics averaged over 24 samples per condition.

| **Metric** | **Traditional**<br>**16 hour**<br>**Vendor A** | **Trinity**<br>**1 hour**<br>**Vendor A** | **Traditional**<br>**16 hour**<br>**Vendor B** | **Trinity**<br>**1 hour**<br>**Vendor B** |
| --- | --- | --- | --- | --- |
| On-Target | 88% | 87% | 93% | 87% |
| Fold-80 | 1.47 | 1.27 | 1.46 | 1.43 |
| %<br>Duplicates | 2.31% | 1.17% | 3.60% | 1.27% |
| SNP F1 | 0.991 | 0.993 | 0.989 | 0.991 |
| Indel F1 | 0.957 | 0.975 | 0.967 | 0.981 |

To evaluate the robustness of the Trinity assay, we sequenced 216 whole exome samples (replicates of HG001 and HG002) with the Trinity workflow and collected metrics. The study utilized 9 flow cells multiplexed at 24 samples per hybridization reaction per flow cell. We evaluated bait sets from two vendors and different hybridization protocols, with multiple replicates for each configuration. Each of the conditions showed metrics within the expected range with low variation across runs (Supplementary Table 1).

We next evaluated Trinity’s compatibility with FFPE and biopsy samples, which are widely used in clinical research despite their challenges of limited DNA quality and quantity. For this, we prepared libraries from 8 gDNA samples from blood, 8 FFPE preserved samples from needle biopsy, and 8 FFPE preserved samples from tissue resection, starting with 50 ng DNA input (DIN ranging from 1.7-4.9 for FFPE and 4.7-8.7 for blood) per sample. The libraries were pooled for hybridization to an exome panel and sequenced following the Trinity protocol.

Trinity showed improvements in mean target coverage due to lower duplicate rates compared to the traditional workflow with the same samples (Supplementary Figure 1). The results were replicated by sequencing 9 of the samples at higher coverage depth.

### Evaluation of different panel sizes

Bait sets targeting the exome and its extensions represent some of the largest panels routinely used in research. Many applications use panels with a more focused target space and therefore a small fraction of the number of baits used for the exome. We explored the compatibility of the Trinity workflow with a variety of such panels to assess performance and determine the amount of starting material required. Testing eight custom panels ranging from 730kb (191 genes, 7816 probes, where ∼.024% of genome is targeted by capture probes) to 43Mb (∼1.39% of genome), we investigated whether (1) adequate target density could be achieved with smaller panels and (2) whether we could predict the required amount of input material based primarily on panel size. Though multiple factors may impact conversion efficiency from an input capturable molecule to a successfully sequenced polony (including panel design, capture efficiencies, and vendor differences), we found that capture performance primarily correlates with panel size relative to genome size. By accounting for this potential capture space of molecules, we found that we could achieve our target flow cell output (∼800M paired-end reads) for a large range of panel sizes by appropriately scaling input into the hybridization reaction. Figure 2A shows the inverse relationship between target size and required input in a log-log plot across a range of commercially available panels (gray circles) and those empirically tested (colored circles). With the eight panels tested, we were able to achieve target density while using total pool input amounts from 0.9 to 24 µg from the largest to the smallest panels, respectively (Supplementary Table 2). The simple relationship between the input amount and the panel size serves as a useful starting point for the loading optimization of any custom panel within the tested range.

**Figure 2.**
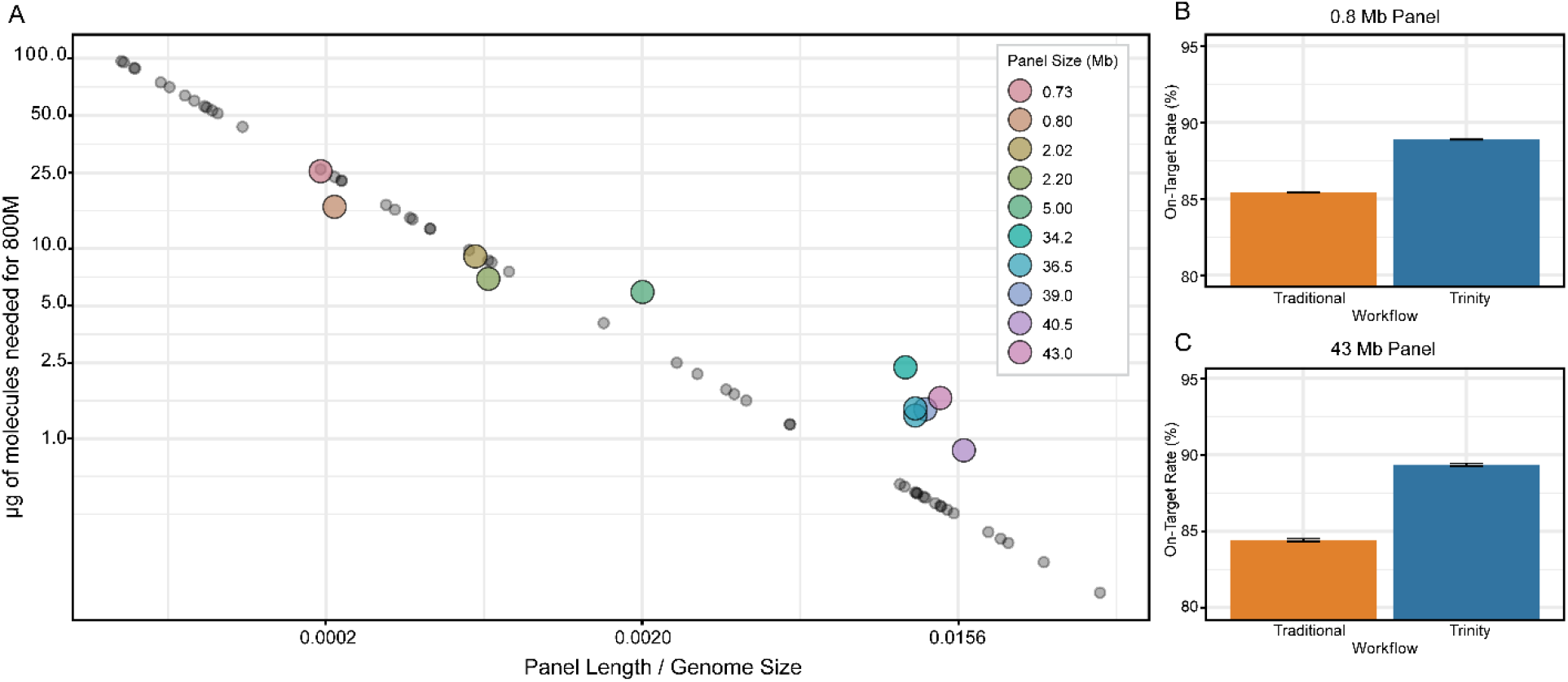
Library loading and performance across panel sizes. (A) relationship between panel size and library loading amount. The DNA amount is cumulative across multiplexed samples, so the left-most point (200 kb panel in human corresponds to ∼96 µg in pool or 1µg per sample in a 96 plex assay). The gray circles provide the approximate loading concentration as a function of panel size for commercially available targeted panels that are similar in size to those that we tested (colored circles). (B) On-target performance for a 0.8Mb pan-cancer panel with Traditional and Trinity workflows (24µg hybridization input, 18µg loading) (C) On-target performance for a 43Mb exome panel (3µg hybridization input, 1.6µg loading).

Importantly, the Trinity workflow maintained high on-target stringency despite these varying hybridization inputs. When testing panels at both extremes of our size range, Trinity demonstrated superior performance compared to the traditional workflow (Figure 2B and 2C). The 800kb panel showed an increase in on-target rate from 85.4% (traditional) to 88.9% (Trinity), while the 43.2Mb panel improved from 84.4% to 89.4%. In addition, mean target coverage, duplicate rate, and indel calling accuracy were comparable or better than in the traditional assay (Supplementary Table 3).

### Evaluation of somatic use cases

We sequenced several somatic panels using the Trinity protocol and 1-hour hybridization to evaluate the somatic use case. A representative example was a 7,901-probe custom oncology panel applied to 10 samples. Compared to the traditional workflow, Trinity showed a nearly two-fold reduction in duplicates (56% to 33%) corresponding to higher library complexity and higher mean target coverage when starting with the same number of input reads (Figure 3A). One of the 10 samples was a reference standard (Horizon myeloid DNA HD829) with 22 known variants at specified allele frequencies. We detected all 22 variants with observed allele frequencies closely matching their expected values (Figure 3B).

**Figure 3.**
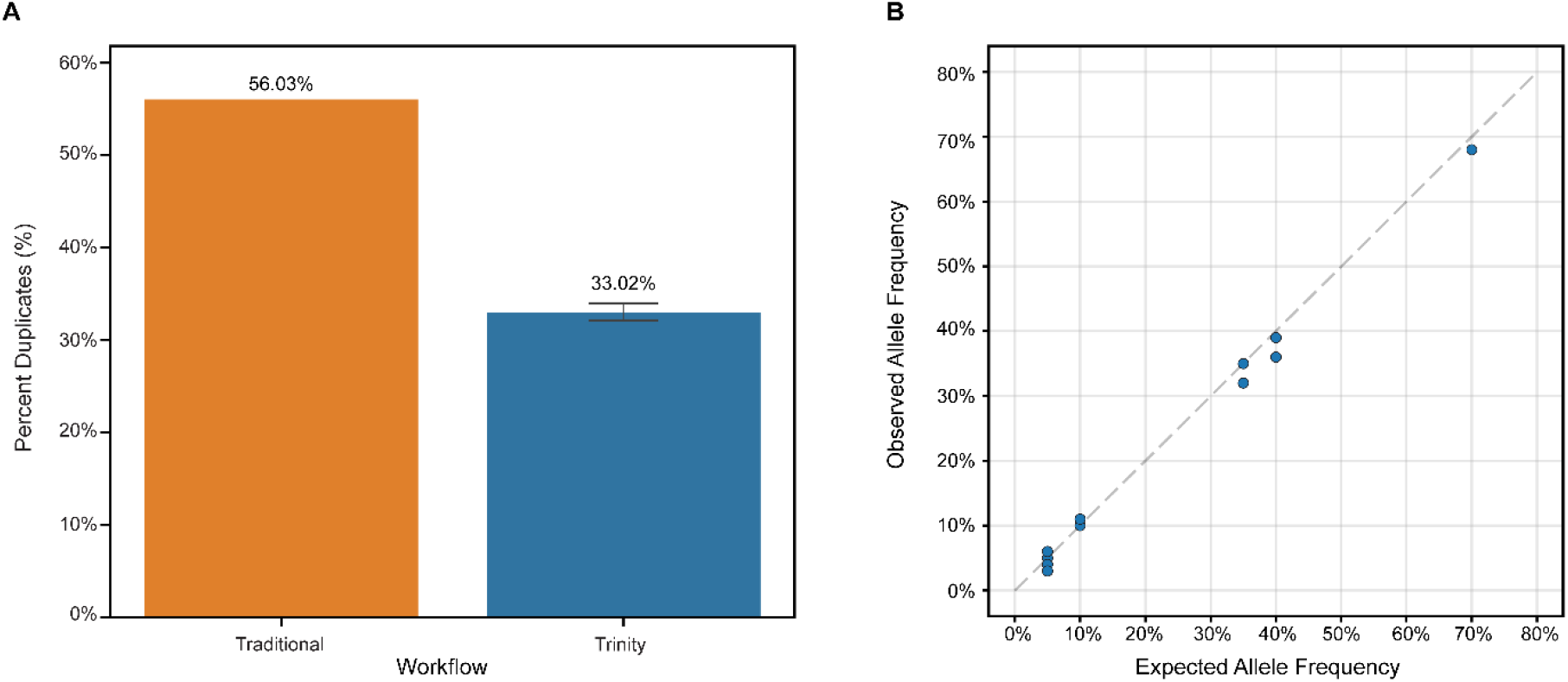
Duplicate percentage and variant calling performance for custom oncology panel. (A) Percent duplicates with Traditional and Trinity workflows. (B) Expected versus observed allele frequency.

### Performance of the PCR-free workflow

The Trinity workflow obviates the need for post hybridization PCR, which presents a unique opportunity to eliminate PCR from all process steps and to evaluate an entirely PCR-free exome. We prepared PCR-free whole genome libraries from the well-characterized Genome in a Bottle samples HG001-HG004, executed the Trinity workflow with exome baits, and sequenced two replicates of each sample on the AVITI sequencer, which employs a rolling circular amplification chemistry, as opposed to a PCR-based approach^23^. As a control, the same samples were processed using the traditional hybrid capture method that employes PCR before and after the hybridization. We also sequenced the same samples using Trinity with PCR prior to hybridization to determine the impact of each PCR step. Using 25 million reads per sample, we saw 1.8% duplicates with the traditional method, 1.3% with Trinity PCR, and 0.41% with Trinity PCR-free. This translates to a 5-fold increase in library complexity when comparing the traditional method to Trinity PCR-free. Figure 4 shows the comparison of variant calling performance among the methods. The PCR-free exome shows a modest improvement in SNP F1-score (from 0.987 to 0.992) but the improvement in indel calling is striking (from 0.938 to 0.985). The number of indel false negative calls is reduced by an average of 67% and the number of indel false positive calls is reduced by an average of 89%. The result demonstrates that most indel errors in exome sequencing are caused by the PCR amplification (as opposed to e.g. coverage non-uniformity). Interestingly, the indel performance gap between Trinity PCR with PCR and Trinity PCR-free is smaller than that gap between traditional and Trinity with PCR, suggesting that more of the benefit comes either from the elimination of the post-hybridization PCR step or from some other aspect of the Trinity assay. To determine the impact of higher coverage on variant calling benchmarking, we repeated the analysis with 80 million reads per sample (Supplementary Figure 2) and demonstrated an indel F1-score above 0.99, approaching the standard set by PCR-free whole genome sequencing^25^.

**Figure 4.**
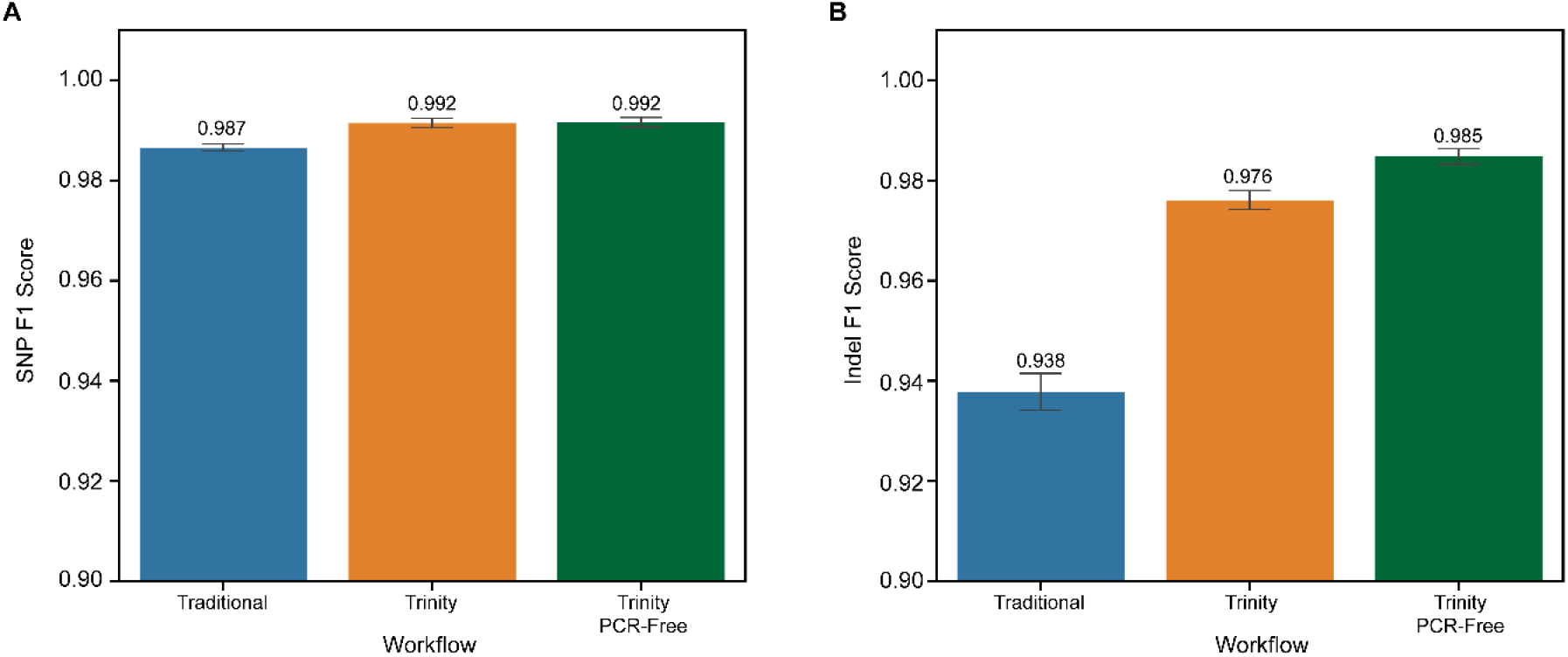
Variant calling benchmarking comparing traditional workflow to Trinity and Trinity PCR-free. (A) The SNP performance is comparable, while the indel F1 scores (B) are markedly higher for the Trinity and Trinity PCR-free assay. Reads were downsampled to 25 million per sample prior to analysis.

We next evaluated the ability of the PCR-free exome data to detect repeat expansions, a variant type reliably called only in PCR-free whole genome data^26^, with initial testing focused on the HTT locus. The HTT (CAG)n repeat expansion is associated with Huntington’s disease and any length over 35 repeat units is pathogenic. We prepared libraries from Coriell samples NA13503 and NA13509, known to harbor a pathogenic expansion in the HTT locus of lengths 45 repeat units and 70 repeat units, respectively. The samples were prepared and sequenced in triplicate using the Trinity PCR-free workflow. Figure 5 shows the allele lengths for both alleles in each sample. The 45-unit expansion is called with the correct length in each replicate. The 70-unit expansion is slightly underestimated, but the reported length is well within the pathogenic range, and the confidence intervals provided by ExpansionHunter contain 70 repeats in each expansion (Supplementary Table 4). The non-expanded alleles in all samples were assigned the expected length. We next looked at 9 additional repeat expansion disease loci across these samples, though we did not expect any of them to harbor pathogenic alleles. We observed sufficient coverage and high confidence length estimates for 6 of the loci, while the other 3 had insufficient coverage for confident calls with the enrichment probe set that we used (Supplementary Figure 3 and 4).

**Figure 5:**
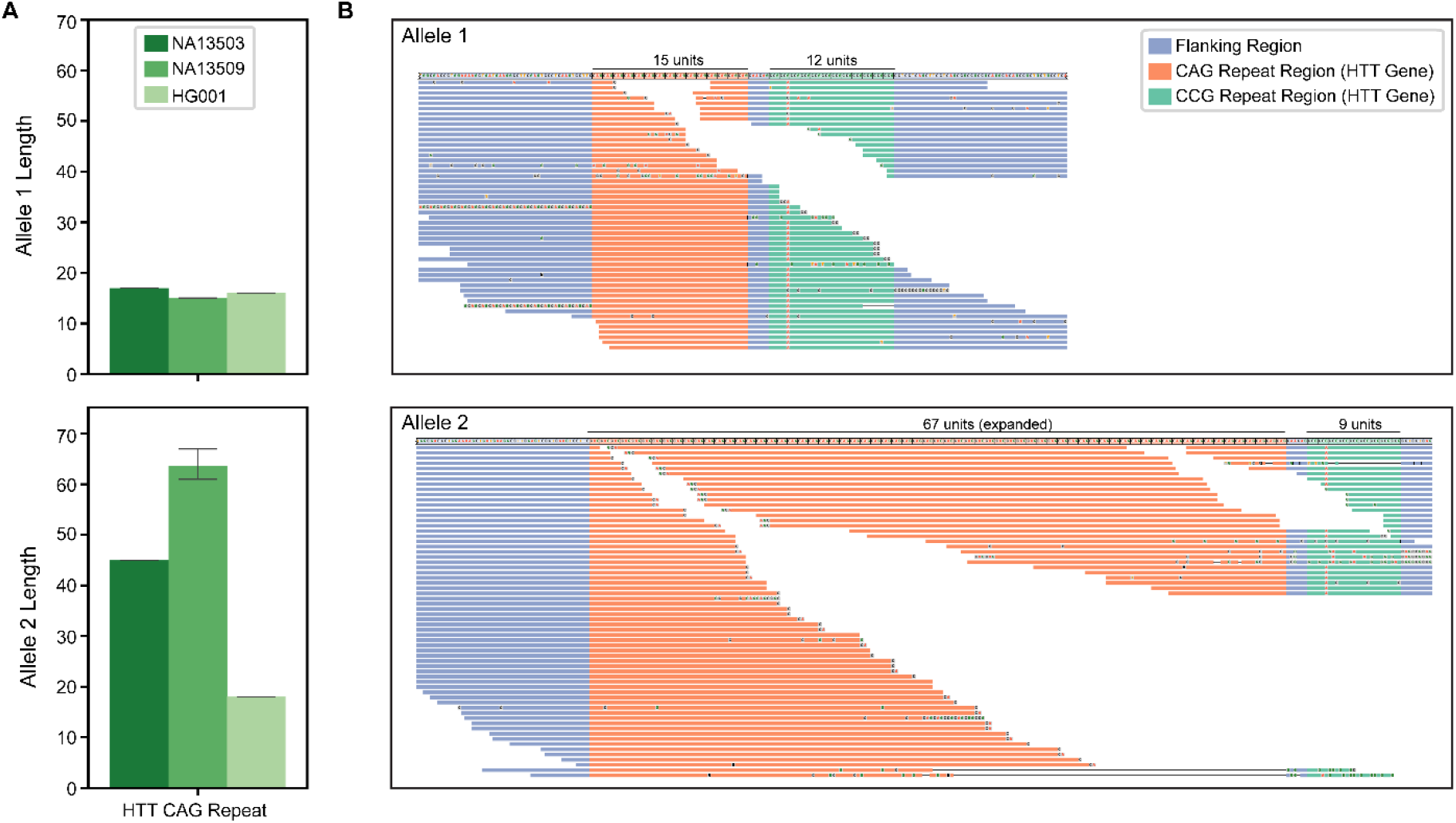
Repeat expansion performance. A: Normal and expanded alleles in the HTT locus based on Trinity PCR-free data. Error bars reflect variation across 3 replicates of each sample. NA13503 and NA13509 both have one expanded allele, while HG001 has two normal alleles. 40M reads were used for each sample. B: Allele visualizations from Reviewer^27^for a representative replicate of NA13509.

## Discussion

We present a simplified hybrid capture workflow that eliminates the steps between hybridization and sequencer loading. Several technological challenges had to be resolved to achieve this simplification without sacrificing specificity. These included the development of a streptavidin flow cell surface that enabled direct capture of the biotinylated probes bound to target, the development of an on-flow cell process for the circularization and amplification of the captured molecules, and the optimization of the hybridization protocol. We also established a relationship between target panel size and DNA input that provides guidance to custom panel users. The workflow was routinely completed within a single work shift enabling users to start a sequencing run on the same day that they began library preparation. The performance of the approach was evaluated by sequencing hundreds of samples and demonstrating high on-target rates, reduced duplicates, and improved indel accuracy. By combining Trinity with a PCR-free library preparation, we enable an entirely PCR-free targeted sequencing assay which further improves indel calling, and shows the potential to detect repeat expansions, a capability which was previously demonstrated only for PCR-free whole genome data.

The current implementation of Trinity has certain limitations that will be addressed in future versions. First, the assay is currently limited to 96-plex unique dual indexing sets. While this is adequate for exome sequencing, where 24 samples typically fill the flow cell, much smaller targeted panels could benefit from a larger set of indices depending on desired coverage. Second, the testing mostly focused on baits from two manufactures (Twist Bioscience and Integrated DNA Technologies). As probe design strategies vary significantly (e.g. single stranded vs double stranded probes or DNA vs RNA probes) additional work is underway to ensure broad compatibility with other manufactures and probe designs. Finally, in the evaluation of repeat expansion calling, we only tested expanded alleles at the HTT locus. For other repeat loci, we evaluated coverage depth in healthy controls but have yet to procure and sequence samples carrying the pathogenic expanded alleles. For loci that were not well covered, we are exploring optimized probe locations.

We believe that additional gains and extensions are possible. For example, given the performance of the 1-hour hybridization, it is likely that the hybridization time could be reduced to as little as 15 minutes without a significant drop in on-target rate for some panel designs and sizes. We are also exploring a modification where the hybridization occurs on the flow cell, thus eliminating additional hands-on steps and equipment from the assay. Finally, by combining low level of the original input libraries with the libraries captured in the hybridization, we can tune the on-target rate for applications that require an enrichment of certain targets along with a uniform genomic background. This combined workflow could be used for imputation or copy number calling. Beyond workflow improvements, the higher library complexity achieved by eliminating post-hybridization PCR opens additional research possibilities. Deep sequencing of these high-complexity libraries may facilitate the discovery of low frequency minor alleles in somatic applications or low abundance species in metagenomics. Work is ongoing to evaluate and optimize these capabilities and to match them to specific applications.

## Methods

### Flow cell functionalization

Passivated streptavidin flow cells were prepared using formulations developed by Element Biosciences, Inc. The flow cells can be purchased as part of the sequencing kits under part numbers 860-00019 and 860-00020.

### Samples and library preparation

Human genomic DNAs from cell line sample HG001, HG002, HG003, and HG004 were obtained from the Coriell Institute. The myeloid gDNA reference sample was purchased from Horizon Discovery (catalog no. HD829). Additionally, representative DNA samples were provided from blood, bone marrow, fresh frozen tissue, FFPE biopsy, and FFPE resection for evaluation.

Extracted DNA samples were fragmented via enzyme treatment or mechanical shearing. Libraries were prepared using the IDT xGen Exome Sequencing Kit Trinity for Element AVITI System (catalog no. 10022463), Twist for Element Exome 2.0 +Comp Library Preparation (catalog no. 109326 or 109327), or Roche KAPA EvoPrep (catalog no. 10154039001) and EvoPlus v2 (catalog no. 09420037001) Library Preparation with xGen Stubby Adapter-UDI’s for Element (catalog no. 10017036), following the vendor’s instructions. For some experiments, a pool of “end-polished” libraries were used. They were prepared with terminal primers and an additional five cycles of PCR prior to library QC. Briefly, 100 ng of each individual-indexed libraries were input into each 50 µl PCR reactions containing 1X IDT xGen HiFi master mix, and 5 µl of xGen Library Amp Primer Mix for Element (catalog no. 10016959). The libraries were amplified in a Thermocycler with the following PCR program: 98°C 45s, followed by 5 cycles of 98°C 45s, 60°C 30s, and 72°C 45s, and a final 72°C extension for 1 minute. The end-polished libraries were purified using 1.0x SPRI beads and quantified by Qubit dsDNA HS Assay Kit (catalog no. 32851).

For PCR-free library preparation, 500 ng gDNA per sample was input into the PCR-free workflow using the Element Elevate Enzymatic Library Prep Kits (catalog no. 830-00009). The final PCR-free libraries were quantified using qPCR as described in the user guide.

### Hybridization and sequencing

Trinity hybridization reactions for IDT and Twist exome panels were carried out according to the Trinity hybridization user guides and using the associated kits (Trinity Sequencing User Guide, IDT xGen Exome Sequencing Kit Trinity for Element AVITI System, Twist for Element Exome 2.0+ Comprehensive Exome Spike-in Library Preparation and Standard Hybridization with Trinity Sequencing Workflow, and Twist for Element Exome 2.0+ Comprehensive Exome Spike-in Library Preparation and Fast Hybridization with Trinity Sequencing Workflow). Additionally, the Trinity workflow with the IDT exome kit was carried out with a one and two-hour fast hybridization (Supplementary Table 5). For this experiment, libraries were pooled by 24-plex for hybridization reactions. After pooling, Human Cot DNA and Trinity Binding Reagent were added to each reaction prior to being dried down as per the xGen Exome Hybridization & Trinity Run Setup protocol. After reactions were dried down, some minor deviations were made from the standard 16-hour hybridization workflow: the pellets were resuspended in 8.5 µL of xGen 2x Hybridization Buffer, 4 µL of xGen Exome v2 Panel, and 4.5µL of nuclease-free water (omitting the xGen Hybridization Buffer Enhancer). These reactions were incubated at room temperature for 5-10 minutes, then sealed and transferred to a thermal cycler, where hybridization would occur with the temperature in user guide.

PCR-free library Trinity hybridization was carried out using a 3 µg PCR-free library pool (quantified by qPCR) as input into the Trinity for Twist fast hybridization workflow followed by the Trinity user guide

Trinity hybridization reactions for non-exome panels from the same vendors were carried out according to the same Trinity protocols with some modifications. For the analysis of the myeloid horizon reference sample (HD829, Horizon Discovery), 6.25 µg of material was hybridized following the Twist for Element Fast Hybridization workflow with 4 µl of the GMS Myeloid gene panel (Genomic Medicine Sweden). For the analysis of the 0.8Mb panel, 24 µg of total end-polished material was hybridized according to the IDT XGen Trinity workflow with 4 µL of the IDT xGen Pan-Cancer Hybridization Panel. For the analysis of the 43Mb exome panel, 5.2 µg of total material was hybridized with the Roche KAPA HyperExome v2 Probes Whole Exome Sequencing solution panel and reagents, plus the Trinity Binding Reagent with reagent volumes and hybridization temperature adjusted to accommodate the addition.

After hybridization was complete, the reactions were removed from the thermal cycler and Trinity run setup proceeded as per the xGen or Twist Exome Hybridization & Trinity Sequencing user guides. The quantity of each diluted hybridization reaction used for final loading was adjusted depending on the library type (stubby or full-length adapter libraries) and hybridization time (1 or16 hours). Trinity hybridization reactions were sequenced on AVITI with a 2×75 or 2×150 Trinity Sequencing Kit, following Trinity Sequencing User Guide.

## Data analysis

FASTQ files were downsampled to a proper read depth, e.g. 30M paired-end reads (30M polonies, 60 M total reads/sample). Reads were aligned to the hg38 or GRCh37 reference genomes using Sentieon BWA-MEM (sentieon-genomics-202308.03) based on target panel coordinates. After alignment, corresponding bed files were applied for panel performance evaluation using Picard (v2.25.0) and Sentieon (sentieon-genomics-202308.03). Variant calling within the defined target regions was performed using DeepVariant (v1.6.1). Variant calling benchmarking utilized hap.py (v0.3.14) against the NIST v4.2.1 truthsets. For the PCR-free data, ExpansionHunter (v5.0.0) was used to estimate sizes of repeat alleles at loci defined within a comprehensive STR catalog corresponding to the hg38 reference. REViewer (v0.2.7) was used to visualize alignment of reads containing tandem repeats.

## Notes

### Competing Interest Statement

The authors are current or former employees of Element Biosciences and may hold stock options in the company.

