## Supplementary Figures and Tables for "A simplified hybrid capture approach retains high specificity and enables PCR-free workflow"

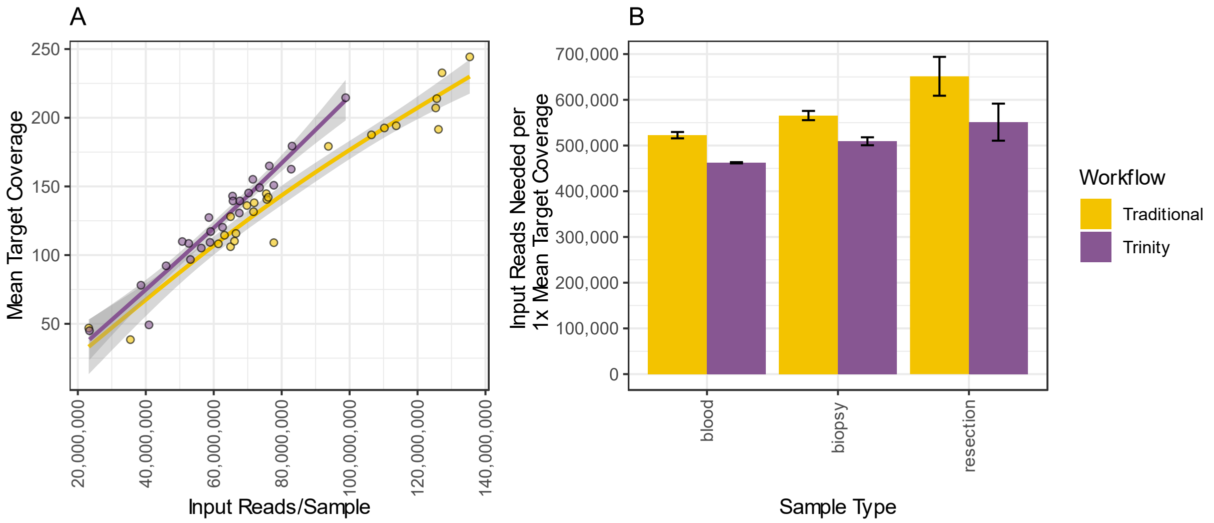


*Supplementary figure 1: Performance on challenging samples. (A) Mean target coverage across samples per workflow (non-downsampled,* n=24 *per workflow). (B) Number of reads needed to generate 1x mean target coverage between workflows, lower values indicate better efficiency in coverage per read sequenced (downsampled to 60 million reads per sample, 8 samples per workflow per sample type).*

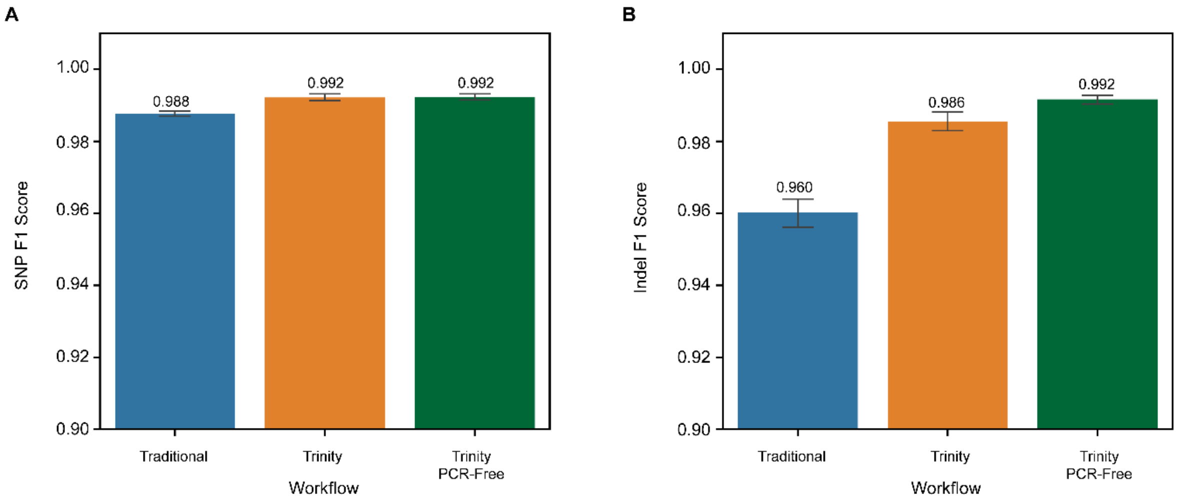


*Supplementary figure 2: Variant calling benchmarking comparing traditional workflow to Trinity and Trinity PCR-free. (A) The SNP performance is comparable, while the indel F1 scores (B) are markedly higher for the Trinity and Trinity PCR-free assay. Analysis performed with 80 million reads per sample).*

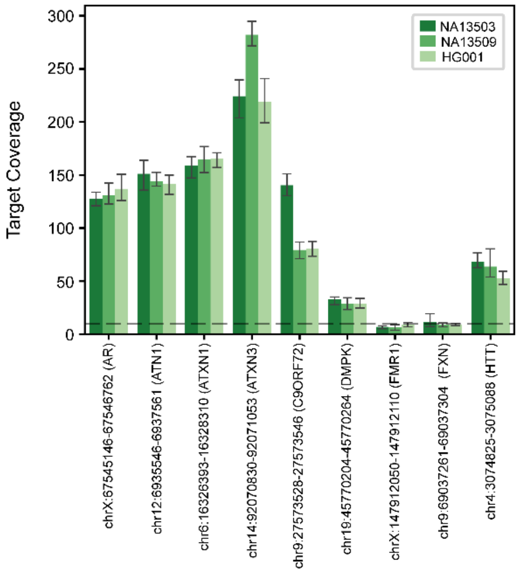


*Supplementary figure 3: Coverage of each repeat expansion locus with the capture probes used. Six of the loci show adequate coverage for confident calls to be made.*

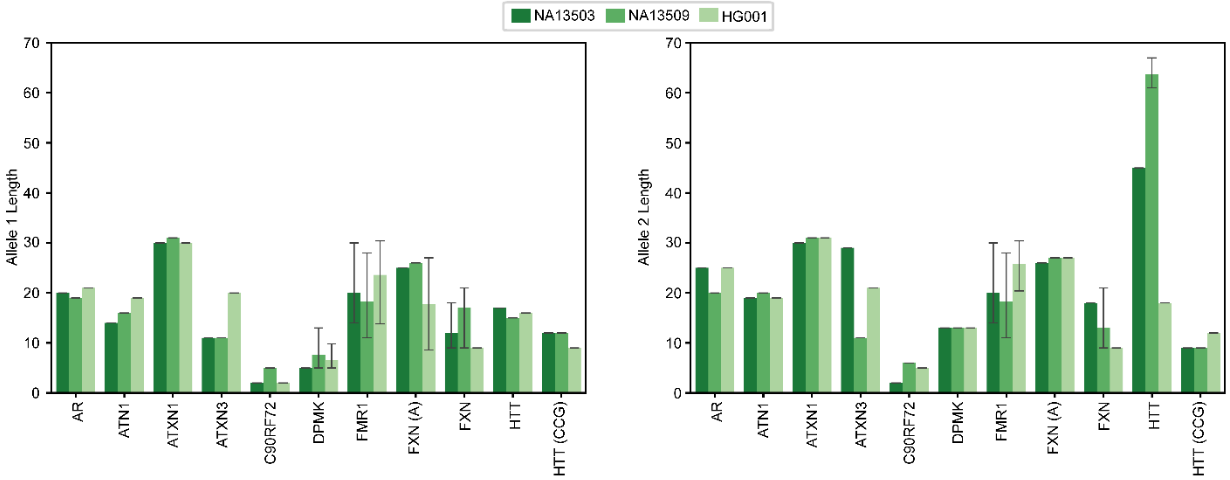


*Supplementary figure 4: Allele lengths at 9 repeat expansion loci. Lengths of alleles in repeat units for 3 samples. Two the samples harbor an expanded allele at the HTT locus and the third sample (HG001) is a normal control. Allele 1 on the left and 2 on the right.*

*Supplementary table 1: Summary of the 216 exome study. Each workflow was evaluated across 3 replicate runs with 24 samples per run. Read lengths were 2x150. Each portion of the table shows a workflow.*

| **Trinity with Vendor A, 16-hour hybridization** | | | | | |
|---|---|---|---|---|---|
| **Run** | **On-target (%)** | **Fold-80** | **Mean Target Coverage** | **SNP F1 Score** | **INDEL F1 Score** |
| **Run1 (Mean)** | 93% | 1.33 | 117x | 0.991 | 0.986 |
| **Run2 (Mean)** | 93% | 1.33 | 121x | 0.991 | 0.985 |
| **Run3 (Mean)** | 93% | 1.35 | 125x | 0.990 | 0.982 |
| **Overall (Mean)** | 93% | 1.33 | 121x | 0.991 | 0.984 |
| **Overall (Coefficient of Variation)** | 0.05% | 0.77% | 3.28% | 0.10% | 0.18% |
| **Trinity with Vendor B, 1-hour Hybridization** | | | | | |
| **Run** | **On-target (%)** | **Fold-80** | **Mean Target Coverage** | **SNP F1 Score** | **INDEL F1 Score** |
| **Run1 (Mean)** | 87% | 1.26 | 95x | 0.992 | 0.978 |
| **Run2 (Mean)** | 88% | 1.26 | 99x | 0.992 | 0.979 |
| **Run3 (Mean)** | 87% | 1.26 | 96x | 0.992 | 0.979 |
| **Overall (Mean)** | 87% | 1.26 | 97x | 0.992 | 0.978 |
| **Overall (Coefficient of Variation)** | 0.20% | 0.12% | 2.08% | 0.00% | 0.03% |
| **Trinity with Vendor B, 16-hour Hybridization** | | | | | |
| **Run** | **On-target (%)** | **Fold-80** | **Mean Target Coverage** | **SNP F1 Score** | **INDEL F1 Score** |
| **Run1 (Mean)** | 86% | 1.35 | 85x | 0.992 | 0.982 |
| **Run2 (Mean)** | 87% | 1.35 | 82x | 0.992 | 0.981 |
| **Run3 (Mean)** | 86% | 1.36 | 84x | 0.993 | 0.981 |
| **Overall (Mean)** | 86% | 1.35 | 84x | 0.992 | 0.982 |
| **Overall (Coefficient of Variation)** | 0.05% | 0.08% | 1.56% | 0.01% | 0.05% |

*Supplementary table 2: Selection of panels tested with Trinity workflow across vendors, and panel sizes. Capture space is Panel Size/Genome size (accounting for differences in species). Hybridization and Loading amounts are total for the pool of samples. Column "µg loading per 800M" is the estimated amount of library needed to load to hit a 800M read target.*

| Vendor | Panel | Panel Size (Mb) | Genome Size (Mb) | Capture Space | Hybridization input (µg) | Hybridization Format | End Polish | Library Loading Amount (µg) | Density (M PF reads) | µg loading per 800M |
|---|---|---|---|---|---|---|---|---|---|---|
| IDT | xGen Pan-Cancer Hyb | 0.8 | 3100 | 0.000258 | 24 | Standard | y | 24 | 1073 | 17.9 |
| N/A | MSK-IMPACT | 2.02 | 3100 | 0.000652 | 18 | Standard | y | 9.9 | 849 | 9.3 |
| IDT | Custom Research Panel | 2.2 | 3100 | 0.000710 | 12 | Standard | n | 12 | 815 | 11.8 |
| IDT | Custom Research Panel | 2.2 | 3100 | 0.000710 | 8 | Standard | y | 8 | 995 | 6.4 |
| Twist | Custom Peanut Panel | 5 | 2556 | 0.001956 | 12 | Standard | n | 8 | 1085 | 5.9 |
| N/A | Sophia Genetics Whole Exome Solution V2 | 34.2 | 3100 | 0.011032 | 6 | Standard | n | 2.7 | 911 | 2.4 |
| Twist | Element Exome 2.0+ Comprehensive | 37.4 | 3100 | 0.012064 | 4 | Fast Hyb | n | 1.5 | 908 | 1.3 |
| Twist | Element Exome 2.0+ Comprehensive | 37.4 | 3100 | 0.012064 | 3.5 | Standard | y | 1.6 | 875 | 1.4 |
| IDT | xGen Exome Hyb Panel V2 | 39 | 3100 | 0.012581 | 12 | Fast Hyb | y | 9 | 925 | 7.8 |
| IDT | xGen Exome Hyb Panel V2 | 39 | 3100 | 0.012581 | 4 | Standard | y | 1.8 | 862 | 1.7 |
| IDT | xGen Exome Hyb Panel V2 | 39 | 3100 | 0.012581 | 4 | Standard | n | 1.8 | 1009 | 1.4 |
| Twist | Alliance Canine Exome | 40.5 | 2500 | 0.016200 | 2 | Fast Hyb | n | 0.94 | 862 | 0.9 |
| Roche | KAPA HyperExome | 43.2 | 3100 | 0.013935 | 3 | Standard | n | 1.5 | 734 | 1.6 |

*Supplementary table 3: Metric comparison of traditional target-capture workflow and Trinity from near-smallest panel tested (B: IDT Pan-Cancer, 0.8 Mb) and largest panel tested (A: Roche Kapa HyperExome, 43.2 Mb). IDT Pan Cancer downsampled to 4M reads/sample, Roche HyperExome downsampled to 60M reads/sample.*

| Panel | Panel Size (Mb) | Workflow | On Target (%) | Duplicates (%) | Mean Target Coverage | Percent Bases Covered at 500x | SNP F1 | Indel F1 |
|---|---|---|---|---|---|---|---|---|
| IDT Pan Cancer | 0.8 | Traditional | 85.43 | 6.69 | 553.93 | 69.72 | 0.999 | 0.858 |
| IDT Pan Cancer | 0.8 | Trinity | 88.88 | 5.99 | 608.26 | 75.65 | 0.998 | 0.931 |

*Supplementary table 4: ExpansionHunter Summary Statistics of the HTT Loci. Repeat ranges defined by gnomAD v4.1.0. Sequences 2x150, analysis 40M reads per sample. Allele lengths in repeat units are classified as follows: Normal ≤ 26, Intermediate 27 - 35, Pathogenic ≥ 36*

| Sample | Loci | Repeat Unit | Classification | Read Support | Number of Repeat Units | Confidence Interval |
|---|---|---|---|---|---|---|
| NA13503_Rep1 | HTT | CAG | Pathogenic | SPANNING/SPANNING | 17/45 | 17-17/45-45 |
| NA13503_Rep2 | HTT | CAG | Pathogenic | SPANNING/SPANNING | 17/45 | 17-17/45-45 |
| NA13503_Rep3 | HTT | CAG | Pathogenic | SPANNING/SPANNING | 17/45 | 17-17/45-45 |
| NA13509_Rep1 | HTT | CAG | Pathogenic | SPANNING/INREPEAT | 15/63 | 15-15/56-78 |
| NA13509_Rep2 | HTT | CAG | Pathogenic | SPANNING/INREPEAT | 15/61 | 15-15/55-76 |
| NA13509_Rep3 | HTT | CAG | Pathogenic | SPANNING/INREPEAT | 15/67 | 15-15/61-83 |
| HG001_Rep1 | HTT | CAG | Normal | SPANNING/SPANNING | 16/18 | 16-16/18-18 |
| HG001_Rep2 | HTT | CAG | Normal | SPANNING/SPANNING | 16/18 | 16-16/18-18 |
| HG001_Rep3 | HTT | CAG | Normal | SPANNING/SPANNING | 16/18 | 16-16/18-18 |
| HG001_Rep4 | HTT | CAG | Normal | SPANNING/SPANNING | 16/18 | 16-16/18-18 |
| HG001_Rep5 | HTT | CAG | Normal | SPANNING/SPANNING | 16/18 | 16-16/18-18 |

*Supplementary table 5: Summary of hybridization input and loading conditions for Trinity workflow using IDT exome kit as a function of hybridization time.*

| Metric | 1-hr Hybridization | 2-hr Hybridization | 16-hr Hybridization |
|---|---|---|---|
| Hybridization input | 15ug | 12ug | 6ug |
| Loading volume | 100ul/200ul | 70ul/200ul | 90ul/100ul |
| Read output (polonies) | 821M | 943M | 899M |
